# Tissue Engineered Axon Tracts Serve as Living Scaffolds to Accelerate Axonal Regeneration and Functional Recovery Following Peripheral Nerve Injury in Rats

**DOI:** 10.1101/654723

**Authors:** Kritika S. Katiyar, Laura A. Struzyna, Joseph P. Morand, Justin C. Burrell, Basak Clements, Franco A. Laimo, Kevin D. Browne, Joachim Kohn, Zarina Ali, Harry C. Ledebur, Douglas H. Smith, D. Kacy Cullen

## Abstract

Although regeneration of damaged axons in peripheral nerves has long been observed, the mechanisms facilitating this growth are not well characterized. Recently, we demonstrated that host axon regeneration could be greatly enhanced by transplanting engineered living axon tracts to guide outgrowth. Here, we used a model of rat sciatic nerve transection to explore potential mechanisms of this facilitated regeneration and its efficacy in comparison with nerve guidance tubes (NGTs) and autografts. Tissue engineered nerve grafts (TENGs) were developed via “stretch-growth” in mechanobioreactors and consisted of centimeter-scale aligned axonal tracts. Either TENGs, NGTs or autografts (reversed nerve) were then transplanted to bridge a 1 cm segmental gap in the sciatic nerve with the mechanisms of axonal regrowth assessed at 2 weeks and the extent of functional recovery assessed at 16 weeks. We observed numerous host axons growing directly along and intertwining with pre-formed axonal tracts in TENGs. This behavior appears to mimic the action of “pioneer” axons in developmental pathfinding by providing living cues for directed and accelerated outgrowth. Indeed, we found that the rates of axon regeneration were 3-4 fold faster than NGTs and equivalent to autografts. It was also observed that infiltration of host Schwann cells – traditional drivers of peripheral axon regeneration – was both accelerated and progressed directly along TENG axonal tracts. These TENG repairs resulted in levels of functional recovery equivalent to autografts, with each being several fold superior to NGT repairs. This new mechanism – which we term “axon-facilitated axon-regeneration” – may be further exploited to enhance axonal regeneration and functional recovery following neurotrauma.

## Introduction

Peripheral nerve injuries (PNIs) present a serious medical concern, with over 100,000 neurosurgical procedures in the U.S. and Europe annually^1–4^. PNI is also a major health concern for warfighters as extremity trauma accounts for as much as 79% of trauma cases in wounded warriors treated in U.S. military facilities, and combat-related blast and/or penetrating injuries generally result in major tissue and peripheral nerve loss^5–8^. Despite the large number of afflicted patients, only 50% achieve good to normal restoration of function following surgical repair – regardless of the repair strategy^9^. This is partly due to insufficient PNI surgical repair strategies that lack biologically-active guidance cues necessary to drive long distance regeneration. Thus, there is a clear need for pro-regenerative “bridges” across segmental nerve defects capable of accelerating axonal regeneration – a major rate-limiting step to the extent of functional recovery.

Following complete nerve transection the axonal segments distal to the injury site rapidly degenerate within hours to days. This followed by a gradual loss of supportive cells (e.g., Schwann cells) that otherwise serve as a natural labeled pathway necessary to guide axon outgrowth to end targets. For long distance axon regeneration, such as down a nerve in the arm, there is a race against time as the slow growth of regenerating axons (approximately 1mm/day) is outpaced by the gradual disappearance of the physical and chemical guidance cues.^10–12^ This commonly results in poor recovery of motor function distal to the original nerve injury^13,14^.

Despite significant efforts, PNI repair strategies have not progressed beyond nerve guidance tubes (NGTs) to bridge small gaps, or autografts – which requires harvesting healthy, uninjured nerve to serve as a living bridge for host axon regeneration – for larger defects, resulting in donor site morbidity as well as other complications^15^. As a result, the field is in need of an alternative technology to promote rapid axonal regeneration across segmental defects while providing a mechanism to attenuate the loss of support cells in the distal nerve segment following major PNI.

We have developed living tissue engineered nerve grafts (TENGs), which are lab-grown nervous tissue comprised of long, aligned axonal tracts spanning two populations of dorsal root ganglia (DRG) neurons. The ability to generate TENGs is based upon the process of axon growth via continuous mechanical tension or “stretch growth^16^. Stretch growth is a natural axonal growth mechanism that can extend axons at rapid rates without the aid of chemical cues, physical guides or growth cones. We routinely replicate this process in custom-built mechanobioreactors through the controlled separation of two integrated neuronal populations (**Fig. 1**). During stretch growth, individual axons gradually coalesce with neighboring axons to form large axonal tracts, or fascicles, taking on a highly organized parallel orientation. TENGs are subsequently created by embedding these living axonal tracts in a three-dimensional (3D) matrix and removing them *en masse* for transplantation^17,18^. This unique platform can generate axons of unprecedented lengths in a very short time frame (5-10 cm in 14-21 days, with no theoretical limit as to the final axon length) from a range of neuronal subtypes and species^16,19–21^.

**Figure 1.**
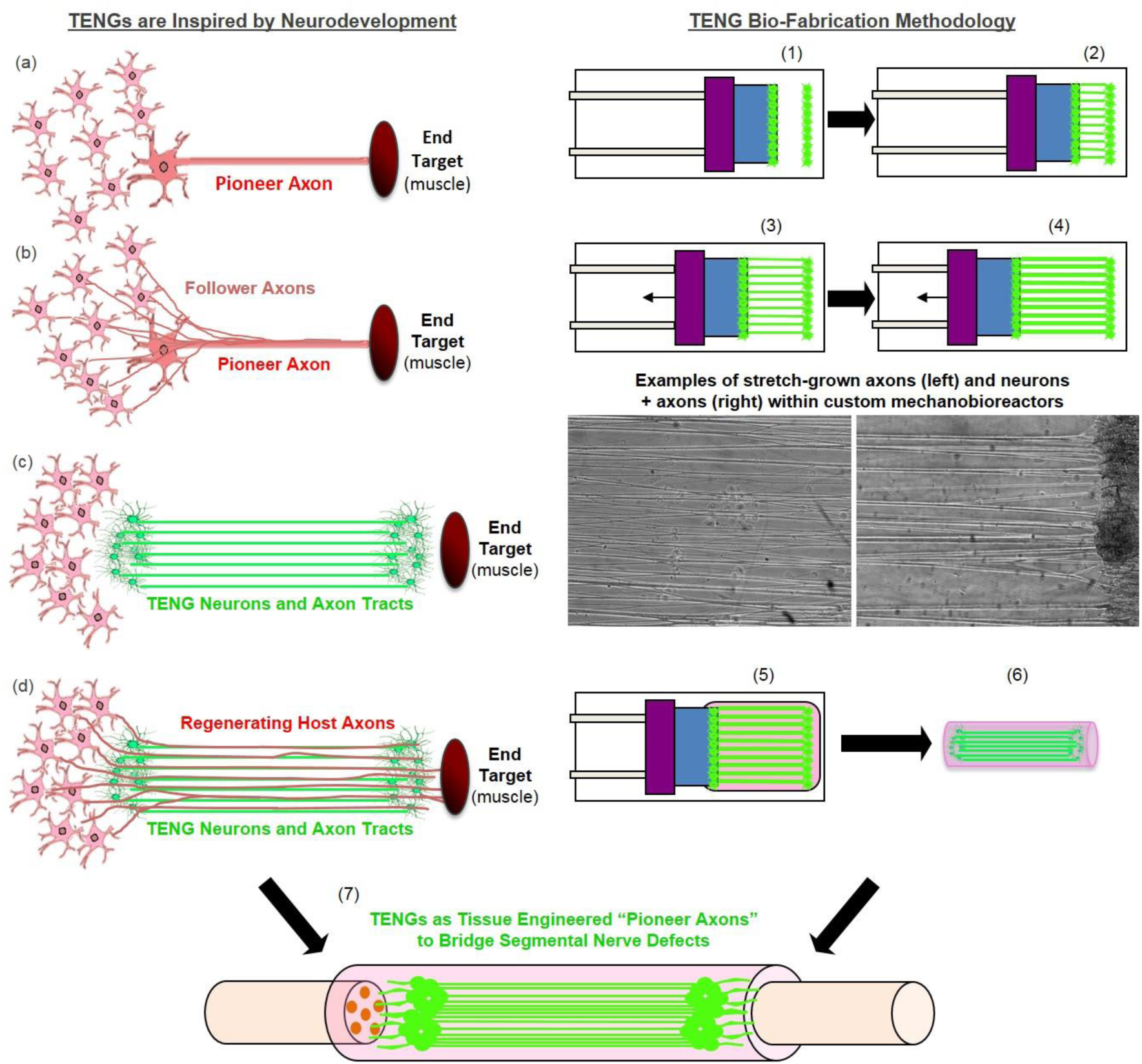
Tissue Engineered Nerve Graft (TENG) Inspiration, Biofabrication, and Surgical Implementation. *LEFT:* TENGs are inspired by axonal pathfinding during nervous system development, where (a) *“pioneer axons”* reach a target first, and then (b) serve as a physical guide for “follower axons” to reach that target. TENG axons are effectively (c) *tissue engineered “pioneer axons”*, thereby functioning as a (d) *living scaffold* to direct and target regenerating host axons across segmental nerve defects. *RIGHT:* TENGs are biofabricated in custom mechanobioreactors via the process of axon “stretch-growth”. Fully formed TENGs – comprised of longitudinally aligned axons encased in a collagenous matrix and rolled into a tubular form – are used to physically bridge segmental defects in peripheral nerve. Briefly, (1) Primary DRG neurons are plated in custom mechanobioreactors. (2) Traditional axon outgrowth integrates two neuron populations. (3) A computer-controlled micro-stepper motor is engaged to gradually separate the two neurons populations, applying mechanical tension to spanning axons. (4) Tension induces axon “stretch-growth”, resulting in increased length, diameter, and fasciculation. “Stretch-growth” occurs for days to weeks at 1-10 mm/day, depending on desired length. (5) Immediately prior to implant, neurons and stretch-grown axons are encased in ECM for stabilization. (6) The ECM containing neurons and stretch-grown axons is “rolled” and transferred into an NGT. (7) NGT containing the cylindrical TENG (neurons/axons embedded in ECM) is then sutured to sciatic nerve to bridge an excised segment.

We have previously transplanted TENGs to study regeneration in a rodent PNI model,^17^ as well as in a rodent spinal cord injury model^22^, with each study demonstrating TENG survival over weeks to months absent any immune suppressive regime. Although these results were promising, for the particular case of PNI repair we did not uncover the mechanism(s) by which TENGs affected axon regeneration, nor did we measure the performance of TENGs compared to the two clincial standards for PNI repair: NGTs and autografts. Therefore, the objective of this study was to investigate the mechanism-of-action (MoA) by which TENGs directly facilitate host axonal regeneration and Schwann cell (SC) infiltration as well as to determine the efficacy of TENGs as compared to standard clinical techniques.

The inspiration for the regenerative MoA of TENGs was based on the observation of axon growth directly along so-called “pioneer” axons during nervous system development. In this case, first, pioneer axons employ pathfinding strategies to find the optimal course to reach and synapse with appropriate targets. Presumably, changes occur on the shaft of the pioneer axons that provide structural cues to direct targeted axon outgrowth from other neurons in the originating site (**Fig. 1**). Thus, we hypothesized that like pioneer axons, TENGs would provide cues to promote host regeneration by direct host axon – TENG axon interactions, ultimately accelerating host axon regeneration across segmental nerve defects and facilitating target reinnervation. We also hypothesized that TENG axons would grow out distally to penetrate into the host nerve, thereby extending the living labeled pathway for regeneration. In the current study, we found that TENGs served as a living scaffold to promote functional restoration at levels surpassing those of NGTs alone and at least equivalent to reverse autografts. Ultimately, tissue engineered “living scaffolds” exploiting potent developmentally inspired mechanisms of regeneration may be useful to facilitate functional recovery following neurotrauma or neurodegenerative disease.

## Results and Discussion

### TENG Biofabrication and Characterization

TENGs were generated within custom-built mechanobioreactors via the controlled separation of integrated neuron populations. DRG explants were isolated from embryonic rats, plated on a stationary membrane and a movable overlapping “towing” membrane, and then virally transduced to express green fluorescent protein (GFP) or mCherry to permit subsequent *in vivo* identification. The two neuronal populations formed connections via axonal extensions across the two membranes over the course of the first 5 days *in vitro* (**Fig. 1**). Then, the towing membrane was slowly pulled away, driven by a precise computer-controlled stepper motor, to physically elongate the axon tracts in micron-scale increments. As previously described, the axon tracts responded to these forces by adding new axon constituents (e.g., cytoskeleton, axolemma, organelles, etc.), increasing diameter, forming fascicles, and lengthening to createlong tracts of living axons (**Fig. 1**)^17–20^. In this manner, axons were stretch-grown to lengths of 1 or 2 cm in length over the next 7 or 14 days *in vitro*, respectively. It is important to note that the stretch-growth media includes specific mitotic inhibitors have been shown to effectively remove the presence of SCs; thus, TENGs are virtually exclusively comprised of DRG neurons and their long axon tracts. Three-dimensional TENGS were then created by encapsulating these stretch-grown living axonal tracts in a collagenous matrix and removing them as a whole for transplantation within a premeasured NGT^17,18^. For implantation, the rat sciatic nerve was exposed and a 1.0 cm or 2.0 cm segment was excised and replaced with an autologous nerve graft (180° reversal of excised nerve), a TENG (within an NGT), or an NGT (either empty or filled with the collagenous matrix used to encapsulate TENGs).

### Acute TENG MoA and Efficacy

To assess the acute regenerative response that occurred in animals treated with TENGs versus autografts or NGTs, at 2 weeks after implantation the nerve/graft zones were harvested and immunohistochemical examination was performed on longitudinal frozen tissue sections. Based on fluorescent reporter expression (i.e. mCherry+ TENG neurons/axons in GFP host rats or GFP+ TENG neurons/axons in wild-type host rats), microscopic examination of tissue sections revealed histological evidence of surviving transplanted DRGs and maintenance of the aligned axonal architecture within all TENG transplants, as expected (**Fig. 2**). To assess host axon regeneration and SC infiltration into the graft zones, sections were immunolabeled for SMI31/32 (neurofilament isoforms broadly expressed by regenerating host axons but not expressed by TENG DRG neurons/axons) and/or S100 protein to denote host SCs. These longitudinal sections provided strong evidence that TENGs were actively facilitating the axonal regeneration process, rather than simply behaving as a permissive substrate for regeneration. In particular, when TENG neurons and axons were placed off-center, host axons altered their growth direction (**Fig. 2c**, white arrow) instead of strictly following the proximal stump trajectory (**Fig. 2c**, gray arrow). This suggests that TENG neurons and axons actively directed and guided the host axon regeneration. Additionally, we employed high-resolution confocal microscopy to enable direct visualization of the mechanisms of axon regeneration at the micro-scale (i.e. visualizing cell-axon and axon-axon interactions). We found regenerating axons had an intrinsic preference to grow directly along TENG axons, thus facilitating vigorous host axon regeneration (**Fig. 2**). We refer to this mechanism as “axon-facilitated axon regeneration” (AFAR), denoting the intrinsic ability of TENG axons to promote host axonal growth directly along their structure.

**Figure 2.**
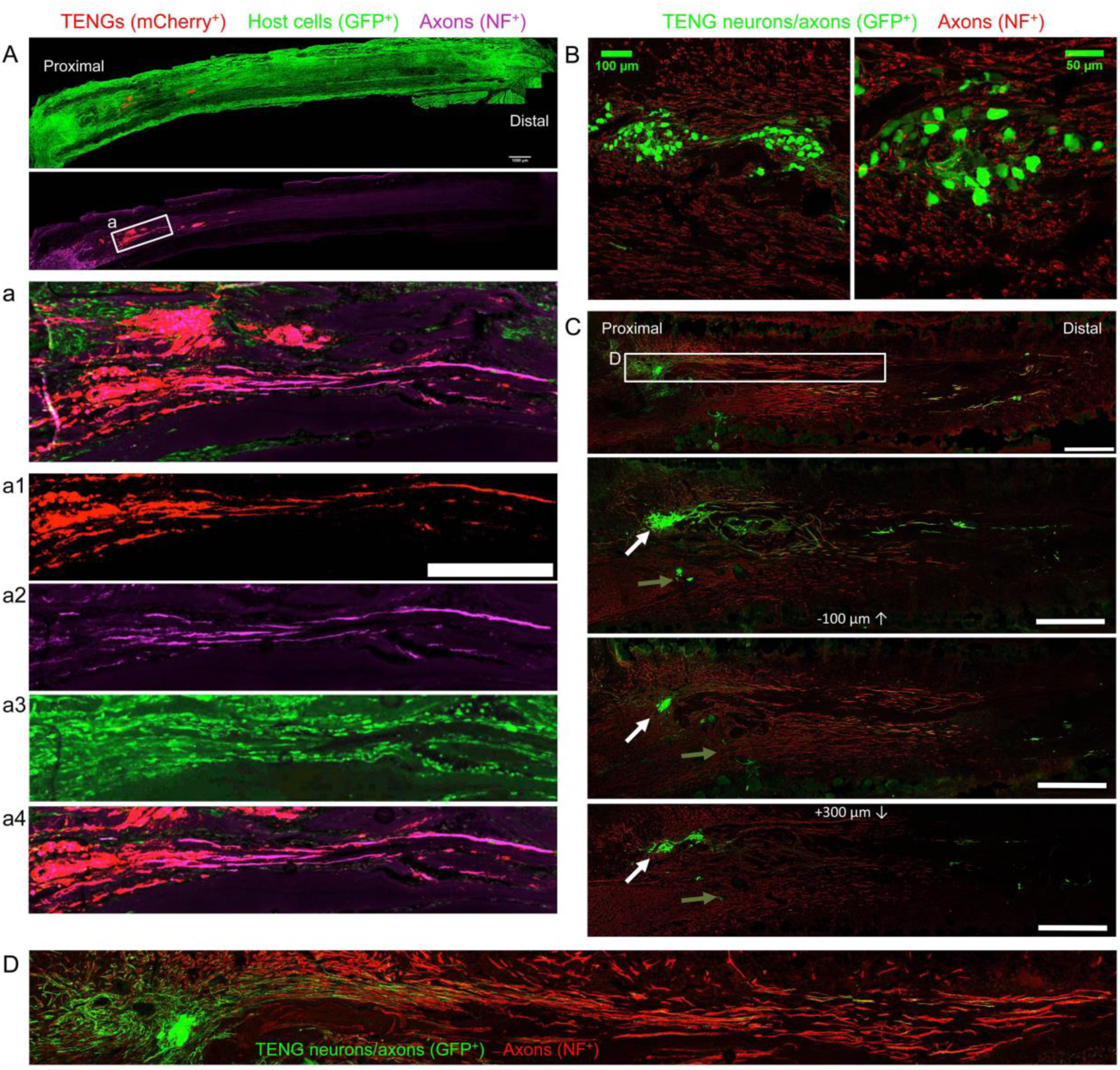
TENG Survival, Maintenance of Architecture, and Mechanism-of-Action following Allogeneic Transplants in Rats. Longitudinal sections across the graft zone at 2 weeks post-implant. (A) Implant of mCherry+ TENGs into GFP+ host rats to discriminate TENG neurons/axons versus host axons, showing robust and directed host axon regeneration directly along TENG axon tracts. (B-D) GFP+ TENG neurons/axons into wild-type host rats. (B) Surviving TENG neurons/axons (green) exhibiting healthy morphology (scale bar: 100 µm). (C) A nerve repaired using a TENG (GFP+) labeled with SMI31 (red) to show axon regeneration. Living TENG neurons and axons were found across the entire nerve graft. TENG neurons/axons were placed off-center to demonstrate that host axons have a preference to follow the path created by the stretch grown axons: the white arrow points out the altered direction of host axon growth through the main cluster of TENG cell bodies and axons; the grey arrow points out the natural axon regeneration trajectory straight out from the proximal stump (scale bar: 1000 µm). This demonstrates that TENG neurons/axons actively direct and guide host axon regeneration. (D) Dense bundles of host axons were intertwined with and grew directly along TENG axons. Collectively, these images provide evidence of a new mechanism of nerve regeneration: axon-facilitated axon regeneration (AFAR) denoted by host axon regeneration directly along tissue engineered axon tracts.

The extent and rates of axonal regeneration across TENGs, NGTs, and autografts were also measured. Host axon penetration across TENGs was greatly increased in comparison to NGTs and was similar to that attained by autografts (**Fig. 3**). In our evaluation of the acute axon regeneration process, we observed two distinct axonal factions: a major bolus of regenerating axons which we termed the “Regenerative Front” (RF), and a much smaller group of more rapidly regenerating axons out ahead of the regenerative front, which we termed “Leading Regenerator” (LR) axons (**Fig. 3**). Notably, the discovery of LR host axons in the distal stump was only found following TENG or autograft repairs, not following NGT repairs (**Fig. 3**). These distinct LR and RF populations of regenerating axons were quantified for all animals at 2 weeks following repair of 1 cm lesions. This analysis determined that TENGs and autografts yielded an equivalent rate of host axon regeneration across the lesion, and that this rate was superior to that observed in animals treated with NGTs (**Fig. 3**). Specifically, TENGs were statistically equivalent to autografts for the RF (p=0.59), with a trend towards TENGs enhancing host LR axon penetration (p=0.081). This was not surprising considering the historical performance of the autograft for small lesions and the fact that the reverse autograft used in this study is a perfect geometric and modality (i.e. ratio of sensory to motor axons) match of the repaired nerve, which is not the case for a human autograft where a sensory nerve is generally used to repair a motor nerve and/or a size mismatch may occur due to limited autograft availability. Based upon the lengths of both the RF and the LRs, TENGs significantly accelerated axonal regeneration versus NGTs (both p<0.01), resulting in axon growth rates 3-to 4-fold faster (peaking at >1 mm/day across the graft zone) than NGTs (**Fig. 3**). We also assessed the density and directionality of axonal regeneration at various locations along the repaired nerve and/or graft. As expected, similar axonal density and morphology was observed across groups in the proximal nerve segment. At the regenerative front, autografts exhibited the greatest longitudinal directionality, while axonal regeneration across TENGs appeared slightly less organized, and NGTs displayed a disorganized webbed morphology, especially at the leading front (**Fig. 3**). Also, NGT axonal density dropped off sharply at the RF, but was more tapered for TENGs and autografts.

**Figure 3.**
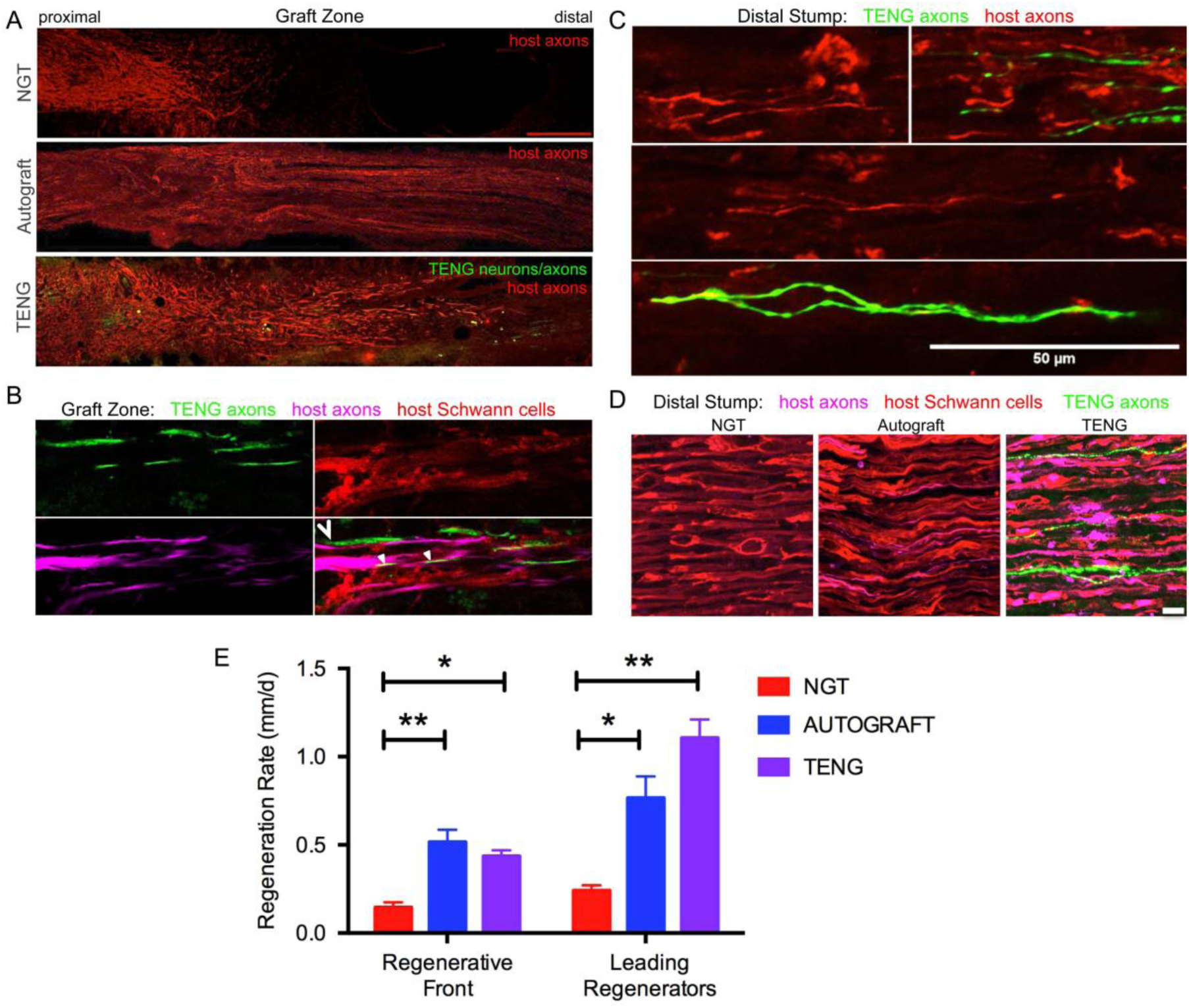
TENGs Accelerate Host Axon Regeneration. Longitudinal sections across the graft zone (A, B) and/or into the distal stump (C, D) at 2 weeks following repair using an NGT, a reverse autograft, or TENG. (A) Axon regeneration as labeled by SMI31 (red, host axons) and GFP (implanted TENG neurons/axons) showing the “regenerative front” – main bolus of regenerating axons – for each repair group. Regeneration through an NGT yielded retarded and disorganized axonal extension. Regeneration through a reverse autograft was highly organized owing to the presence of aligned autologous SCs throughout the graft. Regeneration through TENGs showed host axons following the length of the GFP-labeled TENG axons, with greater penetration and more organization that that found following NGT repair. Scale bar: 1000 µm. (B) Clear discrimination of TENG and host axons, as well as host SCs, across the graft zone. In addition to direct AFAR, host SCs also migrated and organized along TENG axons, creating a “tripartite” regenerative complex of TENG axons: host axons : host SCs. (C) Following TENG repair, a group of host “Leading Regenerator” axons was present in the distal stump – far afield from the “Regenerative Front” – and were seen along with TENG axons that had also penetrated into the distal stump. (D) These accelerated “Leading Regenerator” axons were present following TENG or autograft repair, but were always absent following NGT repair. (E) Plot of quantified axon regeneration measurements (mean ± SEM). Host “Regenerative Front” and “Leading Regenerators” were statistically equivalent following TENGs and autografts repair, and both were superior to NGT repair (*p<0.05, **p<0.01).

Distal to the graft zone, SCs forming the bands of Büngner were found in all groups. However, in the NGT group, there were no axons found in the distal nerve segment, indicating that no axons had crossed the NGTs at 2 weeks post-repair. In contrast, numerous axons were found in the distal nerve structure in animals treated with autografts and TENGs at this early time point (**Fig. 3**). These deeply penetrating host axons within the distal nerve resembled each other in density and morphology for autograft and TENG repairs. Notably, in the TENG group, TENG axons were also found in the distal nerve along with host axons (only host axons were measured as LRs). There were roughly twice as many TENG axons compared to host axons in the distal nerve, demonstrating the permissive environment for axon outgrowth (**Fig. 3**). This unique population of deeply penetrating axons – from both host and TENGs – may extend the living labeled pathway via a combination of neurotrophic support and contact guidance and therefore further promote regeneration. Indeed, these axons may serve to condition the distal environment for the arrival of following axons. Similar to the natural developmental process of the PNS, these axons may prescribe the initial path for subsequent regenerating axons to follow in a process known as “selective fasciculation”^23–25^. Moreover, these axons may serve as regenerative scouts, navigating the more distal environment and sending signals back to the regenerative front in preparation for the unknown territory to come.

As SCs are traditionally believed to be the major drivers of host axonal regeneration across nerve lesions, we also examined the effects of TENGs on host SC infiltration and organization to further understand the effects TENGs had on the active nerve regeneration process. We measured the infiltration/migration distance of SCs from the host tissue, both proximally and distally, into the TENGs and NGTs; such measurements were not warranted for autografts, since they have a homogenous distribution of host SCs from the onset. We found that SC infiltration distance was enhanced in animals treated with TENGs (**Fig. 4**). Conversely, animals that received NGTs showed modest SC penetration from both the proximal and distal ends, with a clear SC-free gap near the center in all cases (**Fig. 4**). Indeed, reconnection of proximal and distal SCs was observed in TENGs but not in NGT groups. In TENGs, the SCs appeared in structures resembling “cables”, potentially precursors to the formation of intra-graft bands of Büngner. Overall, TENGs enhanced SC infiltration into the nerve gap versus NGTs (p<0.01) (**Fig. 4D**).

**Figure 4.**
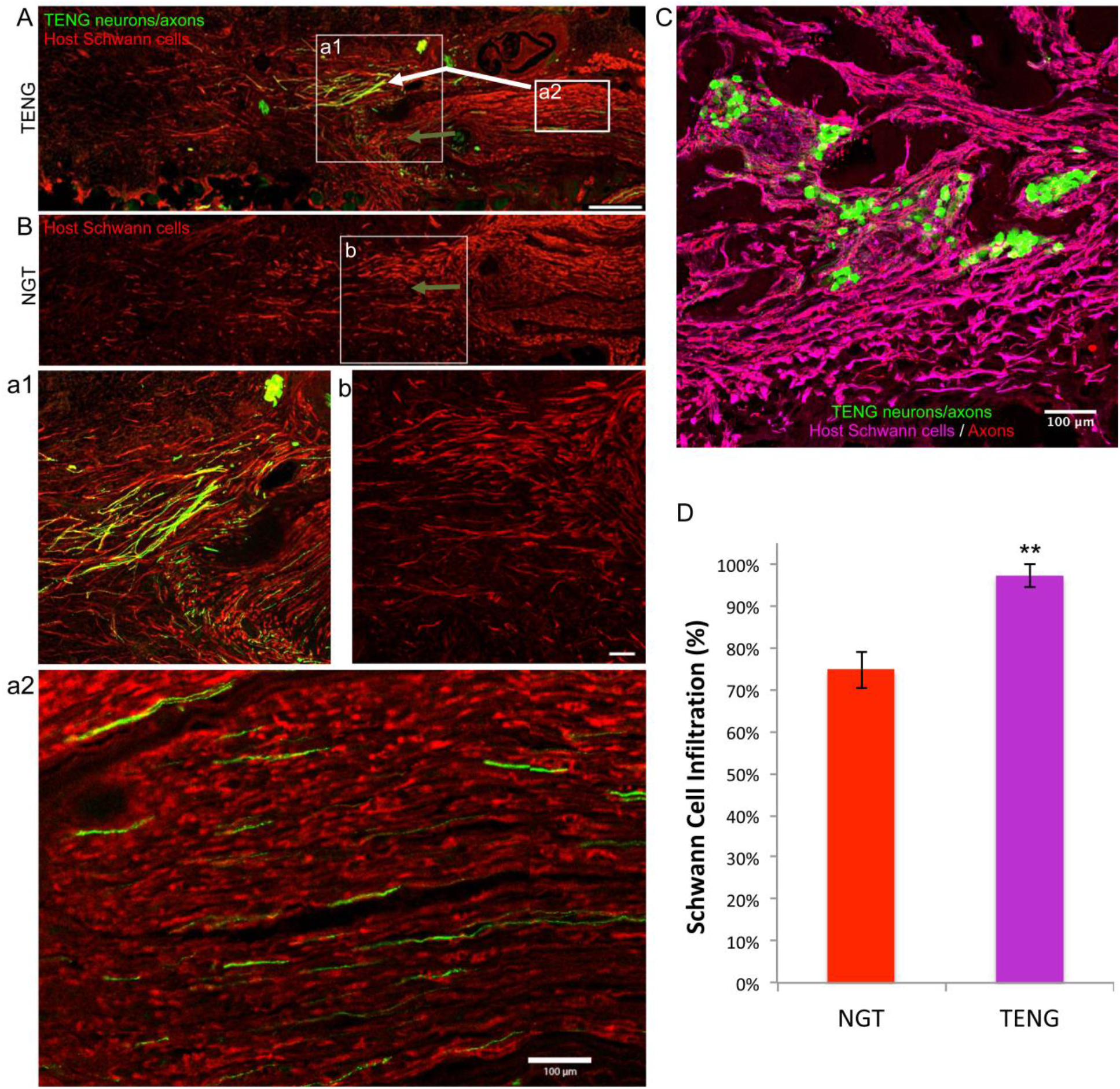
TENGs Direct Host Schwann Cell Infiltration. Longitudinal sections across the graft zone at 2 weeks following repair using TENGs or NGTs. (A, a1) In cases where TENG axons (green) are present off-center (shown in upper half of the boxed section), SC infiltration was markedly directed upward toward TENG axons, indicating that SCs are attracted to TENG axons and alter their migration to infiltrate along TENG axons. (B, b) In NGTs, SCs generally infiltrated linearly from ends, typically observed only in center of NGT (tapered cone). (a2) TENG axons also projected into the distal nerve stump to grow along host SCs. (A, B) Scale bars: 500 µm. (a1, a2, b) Scale bars: 100 µm. (C) Host SCs also directly interacted with TENG neurons; scale bar 100 µm. (D) Plot of quantified SC infiltration measurements (mean ± SEM). TENGs significantly increased infiltration of SCs compared to NGTs (**p<0.01)

Similar to the findings with axon directionality, we observed that SCs had a strong preference to migrate directly to TENG neurons and then directly along TENG axonal tracts. This was likely driven by cell-cell guidance mechanisms, and demonstrated that TENGs actively influenced, directed, and most importanty, acceletated SC infiltration. Taken together, these findings suggest that TENGs play an active role in accelerating the natural regeneration process by encouraging SC infiltration and alignment. This is a critical finding given that following the transection of a nerve, there is a small timeframe during which reinnervation needs to occur for recovery to be complete^26,27^. During this same time, distal axons – which have been physically cut off from their neural cell bodies – undergo a gradual degenerative process and begin to experience a reduction of neurotrophic support leading to aggressive macrophagic degradation^28^. By promoting SC infiltration and organization into aligned columns, TENGs are likely able to promote the synthesis of neurotrophic factors required for regeneration, such as NGF, BDNF and NT-3,^26,28–32^ which would accelerate the reformation of the SC basal lamina^33^. This neurotrophic and structural support would in turn help to encourage axonal sprouts, thereby increasing the likelihood of functional recovery post-repair.

### Chronic Functional Recovery and Nerve Morphometry

We also assessed the degree of functional recovery and mature axonal regeneration at 16 weeks following repair of 1 cm nerve lesions using NGTs, reverse autografts, or TENGs. We found evidence of muscle reinnervation in all animals at this time point based on the presence of compound muscle action potentials (CMAPs); however, the shape and amplitude of the CMAP traces indicated improved muscle health and reinnervation following TENG or autograft repair versus NGT repairs (**Fig. 5A**). The extent of nerve and muscle recovery as measured by CMAP amplitude (normalized to the contralateral muscle) demonstrated a trend towards improvement following TENG or autograft repair versus NGT, but there were not statistically significant differences. Similarly, there was a trend towards improved muscle mass following TENG repair versus NGT, but muscle mass was only significant for autograft versus NGT repairs (p<0.01). In contrast, compound nerve action potential (CNAP) measurements demonstrated a more robust nerve functional recovery following TENG or autograft repairs versus NGT repairs (**Fig. 5C**; each **p<0.01). Nerve morphometry also revealed an increase in the density of regenerated host axons for TENG and autograft repair versus NGTs (**Fig. 5B**; each *p<0.05). Overall, both TENG and autograft repairs elicited superior levels of functional recovery and axon regeneration across multiple metrics as compared to that attained by NGTs, signaling the benefits of endogenous or engineered living scaffolds.

**Figure 5.**
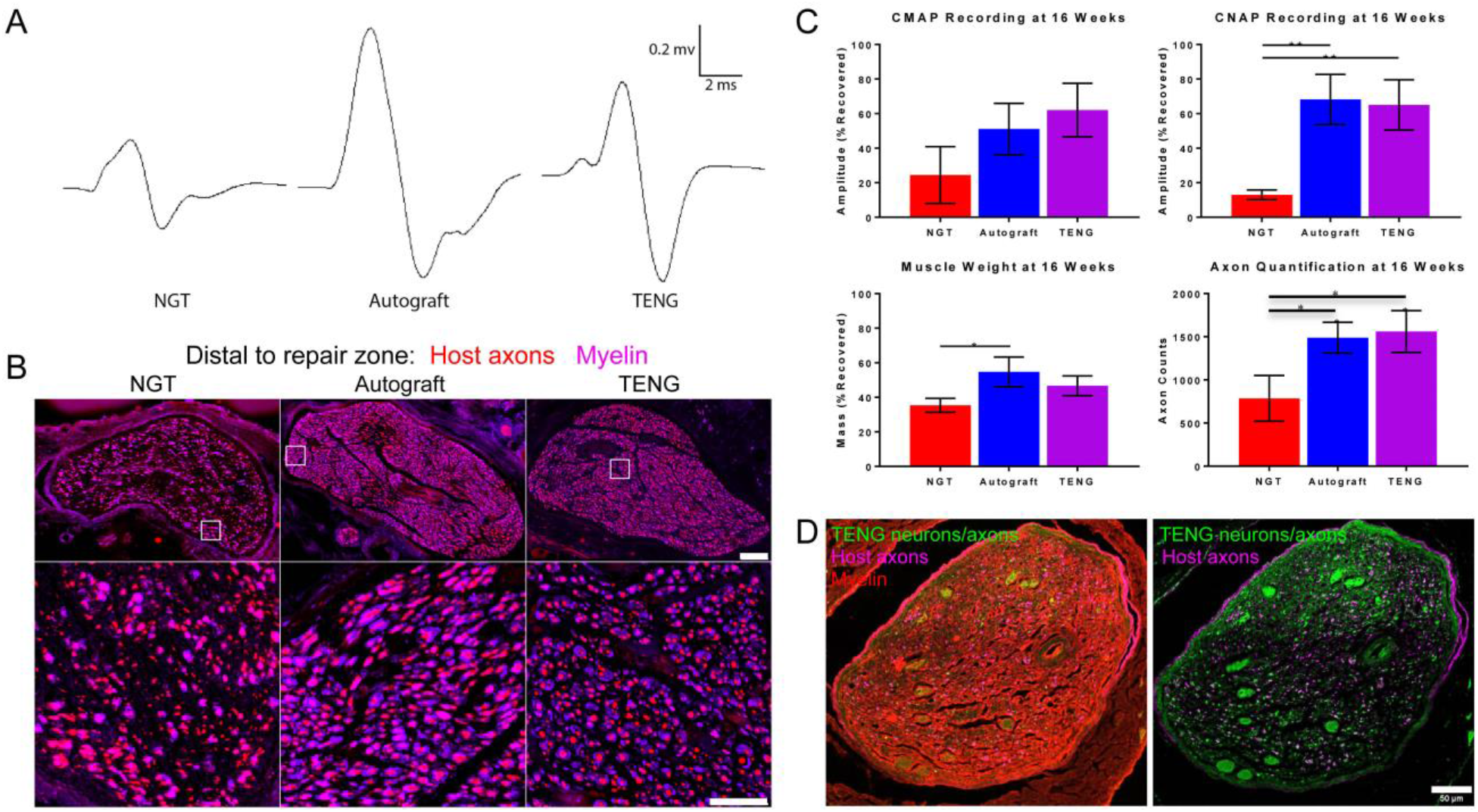
TENGs Facilitate Functional Recovery. (A-C) Functional recovery and structural regeneration at 16 weeks following repair of 1 cm nerve lesions using NGTs, reverse autografts, or TENGs. (A) Representative CMAP traces. (B) Representative nerve morphometry showing nerve cross sections (5 mm distal to repair zone) labeled for axons (red) and myelin (purple). Scale bar (top): 100 µm; scale bar (bottom): 50 µm. (C) Plots of mean recovery levels for CMAP, CNAP, muscle weight, and axon density (mean ± SEM; *p<0.05 and **p<0.01 versus NGT). (D) Chronic nerve morphometry following repair of 2 cm nerve lesions using TENGs. Representative nerve cross sections (5 mm distal to repair zone) at 12 weeks post-repair labeled for axons (purple) and myelin (red), also showing TENG neurons and axons (green). Scale bar: 50 µm.

Finally, we also assessed the ability of TENGs to facilitate axonal regeneration and functional recovery when used to bridge challenging 2 cm defects. Demonstrating proof-of-concept in a longer lesion in rats is a crucial obstacle for a pre-clinical repair strategy. While most engineered bridging solutions are successful for 1 cm lesions due to the strong regenerative capacity of young adult rats, 2 cm defect sizes are generally considered to be beyond the critical defect length in rats and therefore only a subset of effective treatments at shorter (1 cm) defects are effective at longer (2 cm) defects. As we did not know the time course of regeneration and reinnervation for TENGs when used to bridge 2 cm defects, we assessed functional recovery over time using evoked muscle/foot twitch, CMAP, and/or CNAP measurements. This revealed that sciatic nerve stimulation evoked muscle function in some animals as early as 12 weeks following repair with TENGs, with a higher proportion at 16 weeks post-repair. Nerve morphometry analyses supported these functional measures, revealing dense axonal regeneration following TENG repair (**Fig. 5D**). Moreover, TENG axons were also found in the distal nerve out to at least 3-4 months post-repair. These promising results suggest that TENGs may scale to longer lesions and provide regenerative benefits compared to autografts, as TENG axons in the distal nerve may condition and maintain a pro-regenerative environment ahead of regenerating host axons and therefore enable improved functional recovery.

The technique of autologous nerve grafting is considered the “gold standard” and the most reliable choice in repair of major defects in peripheral nerves^34–36^. The introduction of an autologous nerve segment provides physical and biological scaffolding allowing directed guidance of regenerating axons extending to appropriate targets within the periphery. However, complications with the use of autografts exist and are due to the limited supply of donor nerves and the likelihood of donor site morbidity and vulnerability to infection^34–36^. To address the limitations of autografts, alternative strategies to repair damaged nerves have implemented materials of biologic or synthetic origin^13,37–39^. Currently in clinical use are biomaterial-based tubes, such as those comprised of PGA or collagen. These conduits act as a physical guide for axons sprouting from the proximal nerve stump to reach the disconnected nerve segment, which then provide chemical and physical cues to direct regenerating axons to ultimately reinnervate the target tissue^10–12^. However, synthetic conduits have only been clinically successful for the repair of short nerve lesions, and they are typically used for gaps less than 1 cm close to end target^13,14^. Regardless of whether a segmental defect is grafted with a donor nerve or a synthetic conduit, the axons and many supportive cells of the disconnected portion of the nerve ultimately degenerate^10–12^. Due to the relatively slow growth of sprouting axons – as slow as 0.1-0.2 mm/day across an NGT and approximately 1 mm/day across an autograft – as well as the gradual loss of the distal pathway necessary to guide axon outgrowth in the distal segment^10–12^, poor functional recovery of extremities that are far away from nerve damage is seen regardless of the clinically available repair strategy used^13,14^. For example, as is commonly found with brachial nerve injury, elbow flexion may ultimately be regained, but hand function generally is not.^40^

Accordingly, development of alternative means to repair PNI is imperative. Our transplantable, scalable nervous tissue constructs generated via axon “stretch-grown” are a promising strategy to augment and accelerate endogenous regenerative mechanisms following nerve damage. Our findings suggest that TENG axons – via AFAR – mimic the action of “pioneer” axons during development and thereby provide a combination of physical and neurotrophic support to regenerating host axons. Indeed, living axons have been shown to play a critical role during embryonic development in ensuring proper axonal targeting and driving widespread connectivity, and thus may be an essential element of any tissue engineering approach designed to exploit developmental mechanisms in the context of axonal regeneration and targeting. It is important to note that the presence of SCs pre-added to TENGs would likely be detrimental to the mechanism of AFAR, since pioneer axons in development are not myelinated, and the addition of SCs would likely make TENGs highly immunogenic. However, if TENGs were made using autologous cells it would be useful to compare the efficacy of SC-seeded TENGs versus our standard SC-devoid TENGs.

Over the past few decades, alternative tissue engineered solutions have been sought to overcome the limitations associated with autografts and NGTs. These approaches include creating combinations of permissive scaffolds (such as decellularized grafts or hydrogels), extracellular matrix (ECM), trophic factors, and glial or stem cells^41^. Several groups are investigating the use of growth factors and/or Schwann and glial cell combinations to enhance nerve growth within nerve guidance channels^42–51^. Yet others have shown that axons appear to prefer longitudinally-aligned fibers^52,53^ as a regenerative substrate, and attempts are being made to create fibers that elute trophic factors^54^. Unfortunately, these approaches typically ignore the fact that axonal outgrowth *in vivo* occurs along SCs and the basal lamina; thus, strategies that were optimized based on directly promoting axonal outgrowth *in vitro* generally do not mechanistically translate *in vivo*. In addition, these approaches all focus on promoting regeneration of the proximal stump; none of these strategies can delay the eventual degeneration of the distal pathway. Our TENG strategy is an attempt to couple the benefits of these approaches by using tissue engineered axon-based living scaffolds that can potentially provide structural and trophic support not only to serve as a bridge for regenerating axons, but also extend axons into the otherwise axotomized distal nerve segment to maintain the pro-regenerative environment.

Based on the current results, it is evident that TENGs possess a novel MoA compared to NGTs and autografts. TENGs serve as a living scaffold to facilitate nerve regeneration via two complimentaty mechanisms (**Fig. 6**): (1) *axon-facilitated axonal regeneration or AFAR:* TENGs accelerated host axon regeneration directly along TENG axons even in the absense of SCs, and (2) *enhancement of traditional SC-mediated axonal regeneration:* TENG axons increased host SC infiltration and alignment, which in turn accelerated host axon regeneration along these SCs. This newfound form of axon regeneration – AFAR – is unique to TENGs and complements traditional SC-mediated axon regeneration. The MoA of TENGs, which would intrinsically lead to synergistic presentation of neurotrophic, chemotaxic and haptotaxic cues, is only possible with a living scaffold. Notably, no alternative nerve repair approach (NGTs, autografts, acellular allografts) provides living axons to take advantage of the natural AFAR mechanism of regeneration.

**Figure 6.**
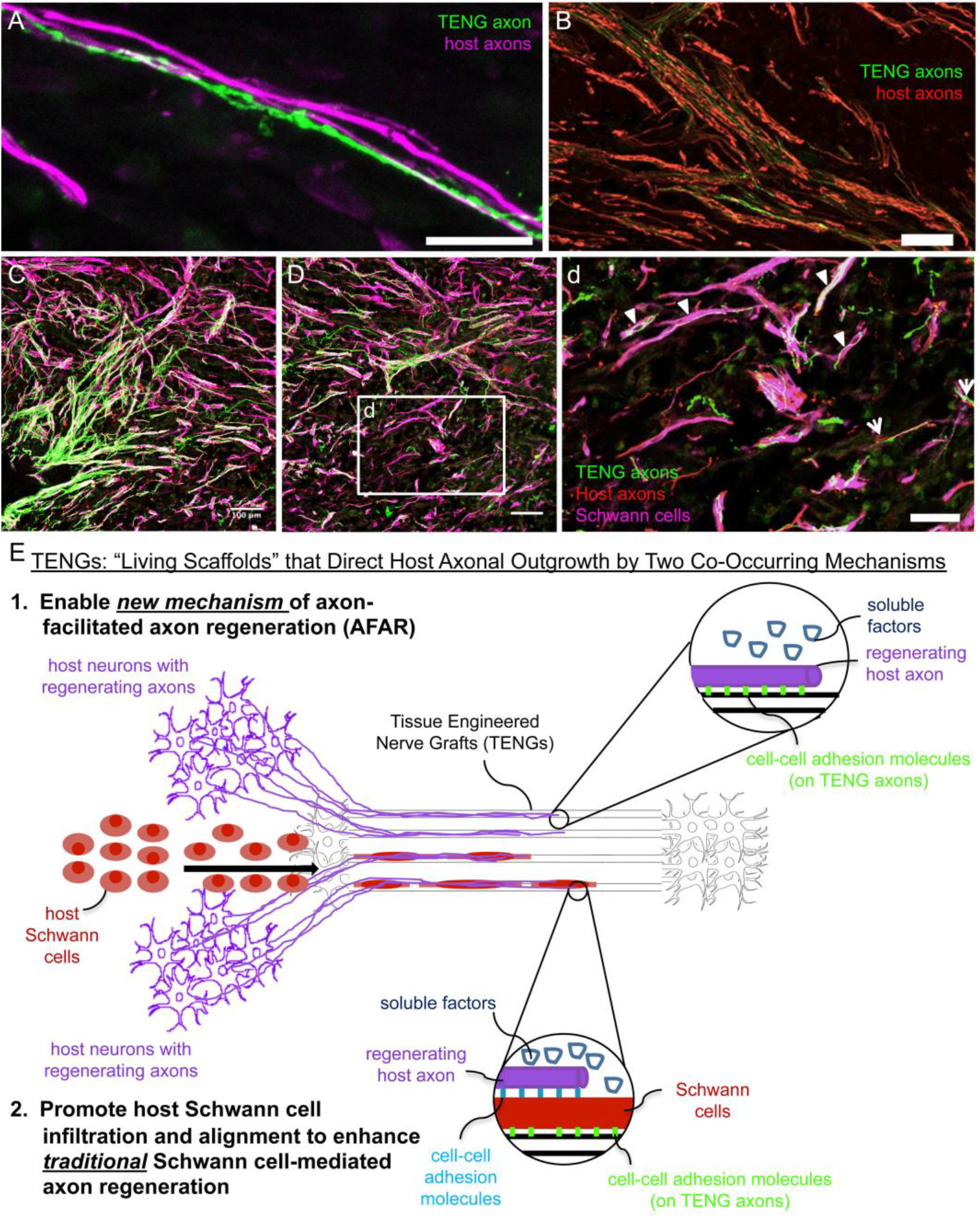
TENGs Mechanisms-of-Action: the Value of Axons. Host axons favor paths created by transplanted TENG axons. TENGs serve as a living scaffold to facilitate regeneration via AFAR on the level of both (A) individual axons (scale bar: 100 µm) and (B) groups of axons (scale bar: 50 µm). (C-D) Tripartite regenerative mechanism: integration of TENG axons with both host axons and host SCs (scale bar: 100 µm). (d) Inset image from (D) showing colocalization of TENG axons with host axons and host SCs. TENGs enhance SC alignment as pointed out by the arrowheads, which then facilitate host axon growth. The arrows show instances of direct AFAR, illustrated by colocalization of TENG and host axons without the presence of host SCs (scale bar: 50 µm). (E) Conceptual schematic depicting the MoA of TENGs, leading to synergistic presentation of neurotrophic, chemotaxic and haptotaxic cues only possible with a “living scaffold”. Overall, TENGS possess novel mechanisms compared to NGTs & autografts: direct AFAR (not possible with autograft), increased SC infiltration and alignment (versus NGT), and robust presence of “leading regenerator” axons (versus NGT). This AFAR – unique to TENGs – compliments traditional SC-mediated axon regeneration.

Overall, we found that AFAR accelerates, directs, and enables robust host axonal regeneration and appears to act in a complimentary manner to traditional SC-mediated axon regeneration. Of note, the mechanism of AFAR does not apply to autografts since the host axons initially within an autograft will inevitably degenerate due to the action of excising the autograft from its original site. This illustrates that TENGs alone have the potential to attain robust axon regeneration without a dependence on proliferating SCs from host tissue, possibly negating the suggestion that SC senescence may lead to a limited amount of axonal regeneration across major nerve injuries regeneration^55^. Additionally, the outgrowth of TENG axons into the distal nerve stump likely enables TENGs to maintain distal nerve segment SCs in a pro-regenerative phenotype and maintain efficacy of the distal pathway to provide a complete guide for regenerating host axons to reach long-distance targets, a mechanism that is also not possible with an autograft.

## Conclusions

Peripheral nerves have a limited capacity for regeneration following injury; however, there is a significant need for next-generation tissue engineered constructs to augment endogenous regenerative mechanisms across segmental nerve defects. To address this need, we have developed living TENGs that consist of aligned axonal tracts routinely generated through the controlled process of axon “stretch-growth”. The concept of TENGs is inspired by developmental neurobiology, where targeted axonal outgrowth occurs directly along pathways blazed by existing axonal tracts, termed “pioneer” axons. When used to bridge segmental sciatic nerve defects in rats, we found that the degree of axonal regeneration and functional recovery attained following TENG repair was statistically equivalent to the “gold standard” autograft repair and consistently superior to NGTs. The unique MoA of TENGs, which facilitates both accelerated axon regeneration across the graft via AFAR as well as sending axons into the distal nerve sheath to prolong its pro-regenerative ability, is the key differentiator from existing technologies and strategies, including the autograft. Moreover, TENGs proved to be superior to NGTs by all three acute metrics measured: host RF axonal regeneration, presence of LR axons in the distal stump, and degree of SC infiltration. In turn, TENGs resulted in several fold increases in the extent of functional recovery following TENG repair versus NGT repair. Our results support the premise that TENGs may enable regeneration across longer nerve lesions following major injury; thus, TENGs may be promising to explore in critical gap (≥5.0 cm) models in a large animal such as swine. Here, TENGs may create a favorable environment for robust and long-distance axon growth that is dependent on the presence of living axon tracts. Overall, this is the first report of a tissue engineered “living scaffold” capable of directly guiding and accelerating axonal re-growth and cell migration across nervous system injuries. This new mechanism of AFAR may be further exploited to enhance target reinnervation and functional recovery following neurotrauma. Therefore, TENGs represent a potentially transformative solution for the repair of complex nerve trauma as well as an option for reconstruction of entire nerve branches following massive tissue loss.

## Materials and Methods

All procedures were approved by the Institutional Animal Care and Use Committees at the University of Pennsylvania and the Michael J. Crescenz Veterans Affairs Medical Center and adhered to the guidelines set forth in the NIH Public Health Service Policy on Humane Care and Use of Laboratory Animals (2015).

### Biofabrication of Tissue Engineered Nerve Grafts

TENGs were generated using dorsal root ganglia (DRG) neurons isolated from embryonic day 16 fetuses from timed-pregnant Sprague-Dawley dames (Charles River). Whole DRG explants were cultured in Neurobasal^©^ medium supplemented with 2% B-27, 500 μM L-glutamine, 1% FBS (Atlanta Biologicals), 2.5 mg/mL glucose (Sigma), 20 ng/mL 2.5S nerve growth factor (BD Biosciences), 20 mM 5FdU (Sigma), and 20 mM uridine (Sigma). Cultures were transduced to express mCherry (AAV1-CB7-CI-mCherry.WPRE.rBG, UPenn Vector Core) or GFP with an AAV viral vector (AAV2/1.hSynapsin.EGFP.WPRE.bGH, UPenn Vector Core).

Explants were plated into mechanical elongation chambers custom-fabricated for stretch-growth. These chambers contained two adjoining membranes of 33C Aclar (SPI supplies) treated with 20 µg/mL Poly-D-lysine (BD Biosciences) and 20 µg/mL laminin (BD Biosciences). One of these membranes, denoted the “towing membrane”, could be precisely moved by a stepper motor. The DRGs were plated in two populations on either side of the membrane interface to allow formation of axonal networks between these two populations. The stepper motor system then separated the populations in micron-size increments until the TENGs reached their desired lengths. Stretched cultures were encapsulated in an extraceullular matrix (ECM) comprised of 3.0 mg/mL rat-tail collagen type I (BD Biosciences) supplemented with 1.0 ug/mL 2.5S nerve growth factor (BD Biosciences). After gelation at 37°C, embedded cultures were manipulated into cylinders, removed from the membranes, and placed within either a premeasured NeuroFlex™(collagen; Stryker), NeuroTube™(polyglycolic acid or PGA; Baxter/Synovis), or tyrosine-derived polycarbonate (TyrPC; Rutgers University) NGT^56^. TyrPC NGTs were synthesized from braided TyrPC fibers (80-100 μm diameter) and dip-coated with a hyaluronan solution (HyStem)^57,58^.

### Peripheral Nerve Surgery and Repair

Experimental subjects were adult male rats (Sprague-Dawley, Charles River). Rats were anesthetized using 2.5% inhaled isoflurane. The left rat sciatic nerve was exposed and a 1.0 cm or 2.0 cm segment was excised and replaced with an autologous nerve graft (1.0 or 2.0 cm long; reversal of excised nerve), an NGT (1.2 cm or 2.2 cm long; filled with the same ECM + NGF used to encapsulate TENGs), or a TENG within an NGT (1.2 cm or 2.2 cm long). There was no difference in the acute regenerative performance of TENGs encased with the PGA (NeuroTube), collagen (NeuroFlex), or TyrPC NGTs, so these groups were combined for statistical purposes. All chronic TENGs were encased in collagen (NeuroFlex) NGTs. Constructs were implanted by inserting the two nerve stumps into the NGT’s ends (1 mm overlap, leaving a gap length of 1.0 cm or 2.0 cm long), which were sutured to the epineurium using four 8-0 absorbable sutures. The wound site was closed with 4-0 prolene or nylon sutures. Experimental groups and group sizes were as follows for sciatic nerve lesion in wild-type rats (unless otherwise indicated), repaired using Autograft (1.0 cm gap: n=8 at 2 weeks, n=4 at 16 weeks), NGT (1.0 cm gap: n=6 at 2 weeks, n=3 at 16 weeks), TENGs (1.0 cm gap: n=6 at 2 weeks into wild-type host, n=3 at 2 weeks into GFP+ host, n=4 at 16 weeks; 2.0 cm gap: n=4 at 12 weeks, n=5 at 16 weeks).

### Functional Assessment

Electrophysiological response was measured at 12 and/or 16 weeks post-transplantation to determine functional regeneration of the nerve following repair. At the terminal time point, animals were re-anesthetized, the surgical site was re-exposed to measure compound nerve action potential (CNAP) and compound muscle action potential (CMAP) recordings.

CMAP recordings were obtained from the tibialis anterior with a bipolar subdermal recording electrode, and a ground electrode (Medtronic, Jacksonville, FL; #8227103) was inserted into the tendon. The nerve was stimulated (biphasic; amplitude: 0–10 mA; duration: 0.2 ms; frequency: 1 Hz) using a handheld bipolar hook electrode (Rochester Electro-Medical, Lutz, FL; #400900) 5 mm proximal to the repair zone. The stimulus intensity was increased to obtain a supramaximal CMAP and averaged over a train of 5 pulses. CMAP recordings were amplified with 100x gain and recorded with 10–10,000 Hz band pass and 60 Hz notch filters.

CNAP recordings were obtained across the graft by stimulating at the proximal site of the sciatic nerve and the distal site on the common peroneal nerve (biphasic; amplitude: 0–1 mA; duration: 0.2 ms; frequency: 1 Hz; 1000x gain, bandpass filter: 10–2000 Hz). The proximal site was stimulated with a handheld bipolar hook electrode (Rochester Electro-Medical, Lutz, FL; #400900) and recorded distal to the repair zone with a bipolar electrode (Medtronic, Jacksonville, FL; #8227410). The ground electrode (Medtronic, Jacksonville, FL; #8227103) was inserted into subcutaneous tissue halfway between the electrodes.

### Nerve Harvest and Histology

At time of harvest, rats were overdosed with sodium pentobarbital. The entire length of the repaired nerves were harvested and placed in 4% paraformaldehyde for 48 hours at 4 °C. The excised tissue was immersed in 30% sucrose solution for 48 hours or until fully saturated. For acute (2 week) animals, the entire repair zone and distal nerve segments were cryosectioned longitudinally (20-25 µm thickness), mounted on glass slides, and immunolabeled using antibodies listed below. For chronic (12 or 16 week) animals, nerve cross-sections were stained and analyzed. Here, the nerve was blocked 5 mm distal to the repair zone and embedded in paraffin. Axial sections (thickness: 8 μm) were taken with a microtome, mounted on glass slides, deparaffinized in xylene and rehydrated with a descending gradient of ethanol. Following rehydration, antigen retrieval was performed in TRIS/EDTA buffer for 8 minutes using a modified pressure cooker/microwave technique. Normal horse serum in Optimax (Biogenex) was applied per manufacturer’s instructions (VectaStain Universal Kit) and sections were incubated at 4 °C with primary antibody (see list below) in Optimax and normal horse serum (VectaStain Universal kit). After washing the sections three times for 5 minutes with PBS/TWEEN, an appropriate fluorescent secondary antibody was applied for 1 hour at room temperature. After rinsing three times, sections were cover slipped. Longitudinal and/or axial sections were labeled using the following primary antibodies (i) SMI31/32 (neurofilament, 1:1500 frozen, 1:1000 paraffin, Covance Research Products), (ii) SMI35 (neurofilament, 1:500, Covance Research Products), (iii) neurofilament-200kDa (1:200, Sigma), (iv) S-100 (1:250, Dako), and/or (v) myelin basic protein (frozen: SMI-94R, 1:500, Covance Research Products; paraffin: CPCA-MBP, 1:1500, Encor). The following secondary fluorophore-conjugated antibodies (AlexaFluor – 488, 568, and/or 647; or Jackson ImmunoResearch) were used as appropriate.

For muscle mass measurements, immediately after euthanasia the ipsilateral and contralateral tibialis anterior muscles were carefully removed cutting the distal tendons, and weighed to obtain the wet muscle mass. Percent recovery was calculated by normalizing the wet muscle mass to the contralateral side.

### Imaging, Quantification, and Statistical Analyses

#### Microscopy

The sections were examined under an epifluorescent microscope (Eclipse E600; Nikon, Melville, NY) and the images were digitally captured (Spot RT Color; Diagnostic Instruments, Sterling Heights, MI). Alternatively, **t**he sections were fluorescently imaged using a laser scanning confocal microscope (AR1, Nikon or LSM 710, Zeiss).

#### Acute Regeneration

For regenerative measurements, sections were captured using the tilescan function in Zen (Zeiss). Regeneration measurements were made by two experienced technicians who were blinded as to the repair group whenever possible, although histological features such as the presence/absence of NGT remnants and presence/absence of TENG neurons/axons made complete blinding impossible in all cases. Acute axon regeneration (at 2 weeks post-repair) was measured across multiple longitudinal sections (minimum of 3 levels) to quantify the distance of the “Regenerative Front” (RF; defined as the main bolus of regenerating axons, always still within the graft zone at 2 weeks following repair of a 1 cm lesion) and “Leading Regenerators” (LRs; defined as the furthest penetrating axons out ahead of the RF axons, generally in the distal stump following autograft or TENG repair, but within the graft zone following NGT repair). RF and LR lengths were measured from the proximal coaptation site (denoted by the sutures in autograft repairs, and the proximal stump transition ∼1 mm in from the edge of the NGT following TENG or NGT repairs). Proximal and distal SC infiltration was also measured from the respective ends of the TENG/NGT to the most distal or proximal S100 staining originating from the respective stump. For images, multiple confocal z-stacks were digitally captured and analyzed. All confocal reconstructions were from full thickness z-stacks from sections 20-25 μm thick.

#### Chronic Morphometry

Axial sections taken distal to the repair zone were labeled for the axonal marker, neurofilament, and myelin (MBP), as described above, to count myelinated axons. The number of axons in each fascicle of each nerve was counted using Fiji by two experienced, blinded technicians. The total axon counts for each repaired nerve was calculated.

#### Functional Measurements

CMAP peak-to-baseline amplitudes were measured and normalized to the contralateral side to calculate percent recovery for each animal. CNAP peak-to-peak amplitude were measured and normalized to the contralateral side to calculate the percent recovery for each animal. Following a normality test, the data was log-transformed as necessary to adjust for non-normality.

#### Statistical Analyses

For all quantitative measures, the group mean, standard deviation, and standard error of the mean were calculated for each group. All quantitative data was analyzed using a one-way ANOVA with repair group as the independent variable and outcome measure (e.g., axon regeneration, SC infiltration, CNAP amplitude, CMAP amplitude, and muscle mass) as the independent variable. When significant differences were detected between groups, Tukey’s post hoc comparisons test was performed for most cases with the exception of the functional data where Dunnet’s multiple comparisons test was performed. For all statistical tests, p<0.05 was required for significance. Statistical testing was performed using GraphPad Prism version 7.03 (GraphPad Software, La Jolla California USA).

## Acknowledgements

Financial support provided by the U.S. Department of Defense [CDMRP/JPC8-CRMRP W81XWH-16-1-0796 (Cullen), MRMC W81XWH-15-1-0466 (Cullen), JWMRP W81XWH-14-1-0100 (Kohn & Cullen), AFIRM W81XWH-08-2-0034 (Kohn, Cullen & Smith)], the Department of Veterans Affairs [BLR&D Merit Review I01-BX003748 (Cullen)], and the National Institutes of Health [NRSA Graduate Research Fellowship F31-NS090476 (Katiyar)].

## Author contributions

D.K.C., Z.A., H.C.L., and D.H.S. conceived of and designed experiments. K.S.K. and L.A.S. carried out TENG biofabrication and *in vitro* imaging/assessment. B.C. and J.K. designed and fabricated nerve guidance conduits. J.C.B., J.P.M., and K.D.B performed lesion and implantation surgeries. J.C.B., L.A.S., J.P.M., and K.S.K. performed electrophysiology assessments. J.P.M., J.C.B, and F.A.L. performed histological evaluations and confocal microscopy. K.S.K, L.A.S., J.C.B, and K.D.B. analyzed data. D.K.C. oversaw all studies and wrote the manuscript. All other authors provided edits and comments to the paper.

## Additional information

Reprints and permissions information is available online at www.nature.com/reprints. Correspondence and requests for materials should be addressed to D.K.C.

